# Localization of the putative recombinase Pf-int to the apicoplast of *Plasmodium falciparum*

**DOI:** 10.1101/2021.10.12.464051

**Authors:** A. V. Berglar, S. S. Vembar, D. N. Gopaul

## Abstract

Diseases caused by apicomplexan parasites, such as malaria and toxoplasmosis cause ∼200 million (worldwide) and 1 million (Europe) infections, respectively, every year. Apicomplexa possess a non-photosynthetic organelle homologous to the plant chloroplast, the so-called apicoplast, that is essential for their growth and survival. This study focused on the Int recombinase, the first protein discovered in *Plasmodium spp*. with the features of a site-specific recombinase, and which has an apicoplast targeting leader sequence at its amino-terminus. Int is conserved amongst several apicomplexan parasites. In the human toxoplasmosis parasite, *Toxoplasma*, Int localizes to the apicoplast and Pf-Int, the *P. falciparum* member, belongs to the group of non-mutable essential genes in *P. falciparum*. A conserved protein that has been shown to be essential at least in one species and that localizes to an essential organelle may become a novel drug target. Therefore, the aim of this study was to confirm the sub-cellular localization of Int in the human malaria parasite *P. falciparum*. Using western blot analysis and immunofluorescence microscopy of *P. falciparum* asexual blood stages, we observed that Int partially co-localized with the apicoplast (to discrete foci adjacent to the nucleus).

## Introduction

Parasites of the phylum apicomplexa cause diseases like malaria and toxoplasmosis and are therefore an important health and socio-economic threat to mankind. The eukaryotic parasite *Plasmodium falciparum* is the main causative agent of Malaria, which generates ∼200 million infections and ∼400,000 deaths every year [1]. *Toxoplasma* gives rise to toxoplasmosis, a severe disabling condition, which is responsible for over one million infections per year in the European region through contaminated food [2]. The lack of effective vaccines highlights the need for novel drug targets against these organisms.

Like all apicomplexan parasites, *Plasmodium* and *Toxoplasma* have evolved from a common photosynthetic red algal ancestor [3]. They harbor an organelle named the apicoplast. The apicoplast was first identified in *T. gondii* [4] and is a non-photosynthetic plastid with four membranes and is homologous to the plant chloroplast. It has been hypothesized to have been evolutionarily derived by secondary endosymbiosis. Even though this plastid has lost the ability to perform photosynthesis over time, it has been retained and serves essential purposes such as fatty acid, isoprenoid precursor, Fe-S cluster and heme biosyntheses[5–7]. The apicoplast carries 1 to 15 copies of a mostly circular 35 kb genome and autonomously performs replication synchronized with schizogony, transcription and translation events [8–10]. These events require plastid DNA copy number management. None of the proteins expressed by the apicoplast genome fulfills this role. The majority of all apicoplast proteins, however, are nuclear-encoded and are targeted to the apicoplast via a N-terminal bipartite targeting sequence, consisting of a translocation sequence and a transit peptide [11–13] (Fig 1). Once imported to the apicoplast, these nuclear-encoded proteins complement the 30 apicoplast-encoded proteins for all remaining plastid activities [14–16].

**Fig 1.**
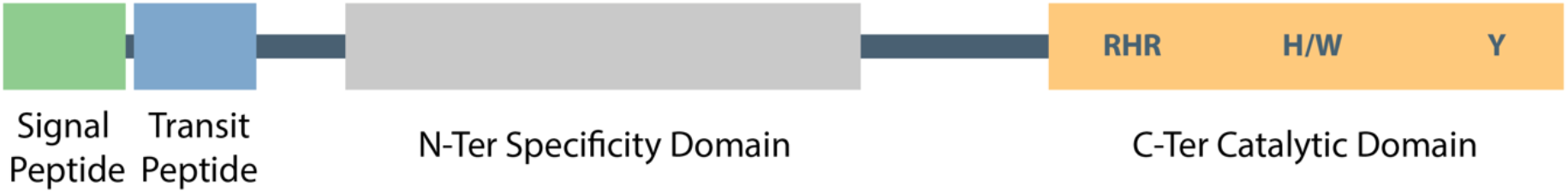
Int protein domain map. Ints as members of Tyr-recombinases are composed of multiple domains. In apicomplexans they also carry a targeting peptide in their N-terminus. The main N-terminal region in Tyr-recombinases are variable in sequence, providing specificity. The C-terminal catalytic domain carries the conserved active site residues involved in DNA cleavage, strand exchange and religation. All members of the Apicomplexa shown here carry the conserved residues of the catalytic domain: R-K/H-H-R-H/W-Y (S1 Fig).

We previously proposed the nuclear-encoded protein Pf-Int (*PF3D7_1308800)* of *Plasmodium falciparum* to be an integrase and we partially characterized it as a putative tyrosine site-specific recombinase (Y-SSR) for DNA binding [17]. Pf-Int has been shown to be essential in a study by Zhang et al. (2018) [18]. It was part of the 2.9% of genes that were not only non-mutable, but that also did not have any Piggy Bac transposon insertions even in their intergenic regions. SSRs in general are involved in a variety of cellular processes such as genome replication, recombination and repair, the impairment of which results in malfunctioning of the organism or interference in the mobility of genetic elements [19–22]. The SSRs require a DNA recombination site ranging between 30 to 200 base pairs in length. These sites contain two motifs with partial inverted repeat (IR) symmetry that flank a central crossover region where DNA strand exchange occurs [23]. Potential outcomes of a recombination event can be insertion, excision or inversion [19] depending on the orientation of the two sites. In practice, this mechanism is used for genome decatenation, partitioning or gene shuffling.

Pf-Int is composed of 490 amino acids (∼58 kDa) and is conserved among six members of *Plasmodium spp*., namely the human infecting *P. falciparum, P. vivax* and *P. knowlesi*, as well as the rodent infecting *P. berghei, P. yoelii* and *P. chabaudi*. Conservation of the C-terminal core part of the protein extends to other members of Apicomplexa in the branches Hematozoa and Coccidia, which are all obligate parasites. Using BLAST, we were able to identify the un-annotated homologs of Pf-Int in *Vitrella brassicaformis* and *Eimeria maxima* (S1 Fig and S2 Fig). This points to the universal distribution of Int among Apicomplaxa that have retained an apicoplast during evolution. Pf-Int shares homology with other well-known Y-SSRs, such as phage λ-Int, P1 phage Cre, bacterial XerC/XerD, the integron encoded VchInt1b (Int4) of *Vibrio cholerae*, and *P. aeruginosa* Int1 mainly in terms of active site residues and polypeptide length of the catalytic domain (amino acid 192-490) [24]. Pf-Int and its homologs in apicomplexa are predicted to encode substantial N-terminal extensions thought to act as plastid-targeting peptides [11]. Nuclear-encoded apicoplast proteins that are targeted into the organelle require multiple trafficking steps from the outermost membrane through the subsequent intermembrane compartments into the lumen [25] (Fig 2). Typically, the upstream portion of the bipartite leader acts as a classical signal peptide that facilitates the co-translational insertion of the protein into the rough endoplasmic reticulum (ER). After cleavage of the signal peptide (SP) by the signal peptidase, the downstream N-terminal transit peptide, similar to those found in plants, is exposed. This transit peptide (TP) directs the trafficking of the protein into the stroma of the apicoplast over either of two different pathways, directly or through the golgi (for details see Fig 2).

**Fig 2.**
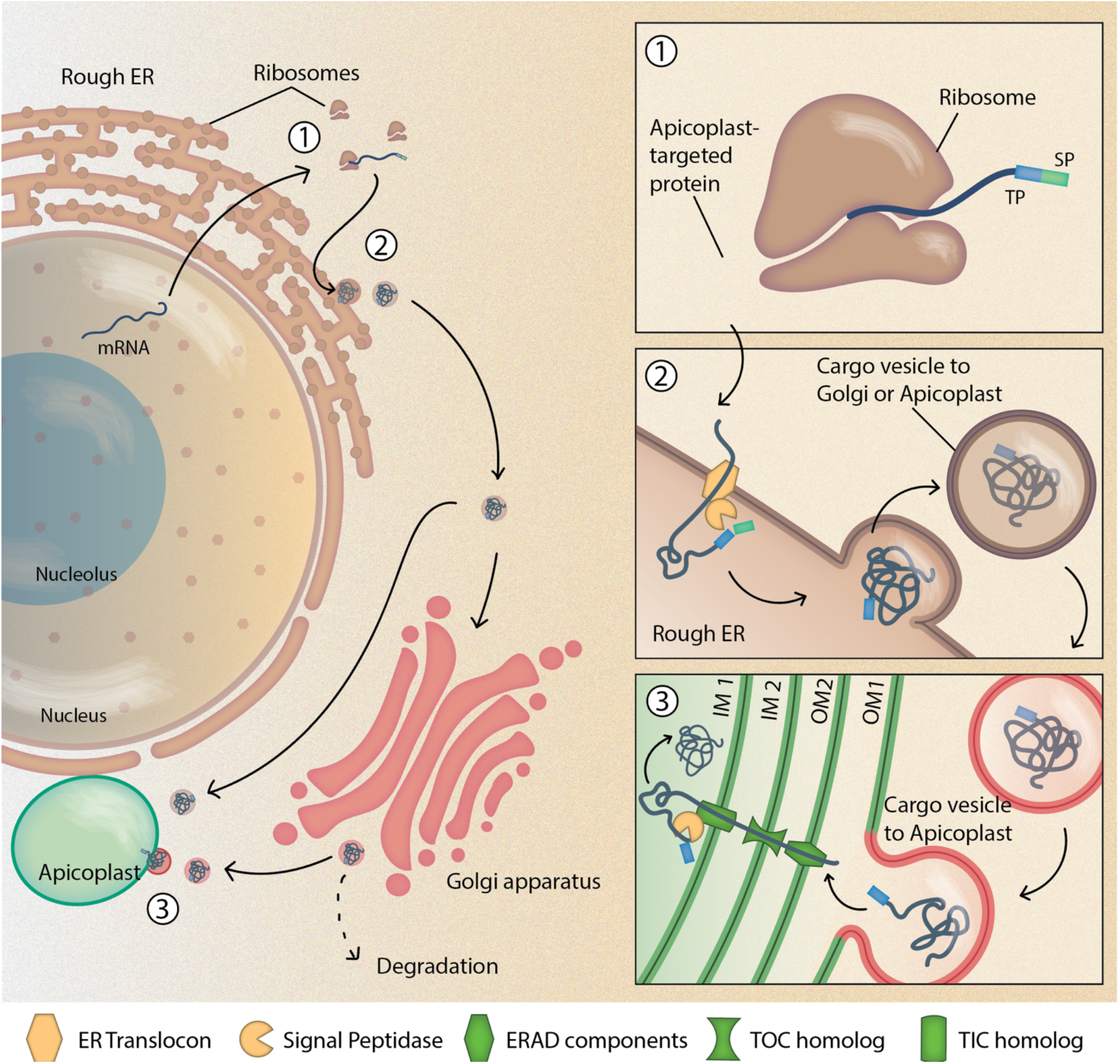
Protein targeting to the apicoplast. 1. The apicoplast destined protein is translated at the ribosome. It has a bipartite leader sequence consisting of a signal peptide (SP, green), which directs the protein to the ER, and a transit peptide (TP, blue), which later mediates the transport across the apicoplast membranes. 2. From the endoplasmic reticulum (ER), where the SP is cleaved off by a peptidase, ER-vesicles carry the cargo either directly to the apicoplast (11), or to the Golgi apparatus, a branchpoint for protein sorting (23,24). 3. Golgi vesicles then fuse with the outermost membrane (OM1) of the apicoplast. Subsequently, the TP guides it across the second outermost membrane (OM2) via an ER-associated degradation (ERAD)-like translocon (25). The second innermost membrane (IM2) is crossed via the outer chloroplast-like membrane (TOC) and the innermost membrane (IM1) via the inner chloroplast-like membrane (TIC) translocons, respectively. In the apicoplast stroma, the TP is cleaved off and the protein adopts its final conformation. Proteins with transmembrane domains anchor themselves in one of the membranes (26,27). Figure © AVB.

The functional role of Int as an integrase is implied with the conservation of critical catalytic residues and protein domains associated with integrases. We hypothesize the role of Int in the apicoplast to be associated with plastid DNA. The homologous protein of Pf-Int in *Toxoplasma gondii* (Tg-Int; *TGME49_*259230, previously TGME49_059230) had independently been identified during a screen of novel apicoplast-resident proteins [26]. The coding sequence of Tg-Int includes a phage integrase domain and a SAP motif. SAP domain proteins are very well conserved from yeast to human, and have been shown to be involved in DNA repair [27], as well as in chromosomal organization [28]. Here, we provide evidence for the co-localization of the integrase to the apicoplast in *P*.*falciparum*.

## Materials and Methods

### Antibodies used in this study

**Table 1.** Antibodies used in this study.

|  | Primary antibodies |  |  | Secondary antibodies |  |  |
| --- | --- | --- | --- | --- | --- | --- |
| Name | Rat Anti Pf-Int C192 | Rat Anti Pf-Int C192 | Anti-HA tag antibody [12CA5] (ab1424) | Amersham ECL Rat IgG, HRP-linked whole antibody (from goat) | Goat anti-Rat IgG (H+L) Cross-Adsorbed Secondary Antibody, Alexa Fluor 488 | Goat anti-Mouse IgG (H+L) Cross-Adsorbed Secondary Antibody, Alexa Fluor 568 |
| Clonality | Polyclonal | Polyclonal | Monoclonal | n/a | Polyclonal | Polyclonal |
| Host species | Rat | Rat | Mouse | Goat | Goat | Goat |
| Commercial supplier or source laboratory | Eurogentech | Eurogentech | AbCam | Amersham (GE Healthcare) | ThermoFisher Scientific | ThermoFisher Scientific |
| Catalogue or clone number | n/a custom-synthesized | n/a custom-synthesized | ab1424 | NA935 | A-11006 | A-11004 |
| Batch number | n/a | n/a | n/a | n/a | n/a | n/a |
| Antigen(s) used to raise the antibody | Purified recombinant Pf-Int C192 | Purified recombinant Pf-Int C192 | n/a | n/a | n/a | n/a |
| Public identifier from the Antibody Registry | n/a | n/a | AB_301017 | AB_772207 | AB_2534074 | AB_2534072 |
| Final antibody concentration or dilution | 1:500 (WB) | 1:250 (IFA) | 1:500 (IFA) | 1:10,000 (WB) | 1:1000 (IFA) | 1:1000 (IFA) |
| See figure | 3A | 3B, 3C | 3C | 3A | 3B, 3C | 3C |

#### P. falciparum parasite culture

Red blood cells were obtained from the Etablissement Français du Sang of Necker hospital, Paris, under agreement with Institut Pasteur, and following the guidelines for informed consent of donors for the use of blood or its derivatives for research purposes. The red blood cells were completely de-identified before researchers accessed the samples. *P. falciparum* blood stage parasites from the 3D7 strain [29] were cultured using modifications to the method described by Trager and Jensen [30]. Parasites were grown in 0^+^ human erythrocytes in RPMI 1640 medium containing l-glutamine (Invitrogen) supplemented with 2X Hypoxanthine, 50 μg/ml gentamycine, 5% (v/v) human serum (PAA Laboratories GmbH) and 5 % v/v Albumax II (Invitrogen) at 37°C in a gas environment of 5 % CO_2_, 5 % O_2_ and 90 % N_2_ in Falcon culture flasks. The culture medium was changed every 48 h by aspiration and the parasite were diluted in fresh RBCs according to parasitemia. Synchronization of cultures consisted of two consecutive 5% sorbitol (Sigma) treatments [31]. We estimated by giemsa staining that the parasites were synchronized within a window of ∼6 h.

#### Fractionation of P. falciparum total protein content

The total protein content of synchronized *P. falciparum* parasites was fractionated into cytoplasmic and nuclear fractions after Oehring et al. (2012) [32] with some modifications. Briefly, parasites 2×10^10^ ring stages (2-20 hpi); 10^10^ trophozoites (20-34 hpi); 5×10^9^ schizonts (34-48 hpi) were released from RBCs by saponin lysis and washed three times in PBS. Parasites were lysed in a hypotonic cytoplasmic lysis buffer CLB (20 mM HEPES (pH 7.9), 10 mM KCl, 1 mM EDTA, 1 mM EGTA, 0.65 % NP-40, 1 mM DTT, Complete TM protease inhibitors (Roche Diagnostics)) for 5 min on ice. Nuclei were pelleted at 2,000 x g for 5 min and the supernatant saved (cytoplasmic extract; fraction 1). After four to seven washes in CLB the pellet was solubilized in 1.5 ml (3 ml) SDS extraction buffer SEB (2 % SDS, 10 mM Tris-HCl (pH 7.5)) for 20 min under constant agitation at room temperature, cleared by centrifugation at 13,000 rpm for 15 min and saved (SDS extract; fraction 2). Large volumes for SDS-PAGE were concentrated by trichloracetic acid (TCA) precipitation.

### TCA precipitation

One volume of TCA stock (100% (w/v) TCA) was added to four volumes of protein sample and incubated for 10 min at 4°C. The samples were spun in a microcentrifuge at 14,000 rpm for 5 min. The protein pellet was washed twice with 200 μl cold acetone (Carlo Erba Reagents). The pellet was dried by placing the tube in a 95°C heat block for 5-10 min. For SDS-PAGE, sample buffer was added and the samples were heated for 10 min at 95°C before loading onto the gel.

### Western blot analysis of Pf-int expression in blood stage parasites

An SDS-PAGE with fractionated protein samples was run in 6x SDS Loading buffer (Laemmli, 2% SDS) to a final concentration of 1x. Samples were run on a 4-12% Bis-Tris SDS gel (NuPage, Invitrogen), 40 min at 200 V in MES running buffer (Novex® Invitrogen) or 60 min at 180 V in MOPS (Novex® Invitrogen) running buffer. As molecular marker, we used the PAGE RulerTM Prestained Protein ladder 10-180 kDa (Fermentas). Recombinant and purified Pf-int was used as a control. Its preparation is described elsewhere [17].

Western blot analyses were carried out using a rat antibody raised against purified recombinant Pf-int (aa 192-490) (Eurogentec, Belgium). The protein content was transferred to a nitrocellulose membrane using the Trans Blot Turbo (BioRad) device program MIXED MW (5-150 kDa, 7 min, 1.3 A up to 25 V). The membrane was immediately placed into saturation buffer and incubated for 1 hour at RT with agitation, then incubated with the primary antibody at 1:500 dilution for 1 h at RT or o/n at 4°C. The membrane was then incubated with the secondary antibody (1:1000 dilution of goat αrat-HRP) for 1 h at RT or o/n at 4°C. 4x 5 min Washing buffer washes at RT and with agitation were done after each incubation step. Blots were developed using the femto kit (Thermo Fisher Scientific).

### Localization studies in *P. falciparum* via Immunofluorescence assay

Localization studies were carried out using a rat antibody raised against purified recombinant Pf-int (aa 192-490) (Eurogentec, Belgium). Fixed parasites were spotted in each well of the microscopy slide and air-dried at RT in order to allow the parasites to adhere. The wells were blocked for 30 min at RT with 15 μl PBS+1% BSA (filter sterilized, Sigma Aldrich). The primary antibody solution was prepared in PBS + 1% BSA. The optimal antibody concentration was determined by dilution series. No primary antibody or pre-immune serum controls were used. 10 μl of primary antibody solution were added to the corresponding wells and incubated in a humid chamber for 1 h at RT or o/n at 4°C. Each well was washed thrice with PBS for 5 min at RT. 10 μl of the secondary antibody (AlexaFluor®), prepared in PBS + 1% BSA, were added to each well at a dilution of 1:1000 and incubated for 30 min at RT. Each well was washed thrice with PBS + 0.5% Tween-20 for 5 min at RT. After the final wash, 3 μl of PROLONG Gold Antifade + DAPI solution were added to each well and the slide was covered with a clean coverslip (Menzel) and sealed with nail varnish. The finished slide was visualized under an Eclipse 80i microscope (Nikon).

## Results

In this study, we performed immunolocalization studies of a putative apicomplexan integrase Int using immunofluorescence assays (IFA) in *P. falciparum*. In the following, we report the results of our investigation.

### *P. falciparum* Int is translocated to the apicoplast

To determine the localization of Pf-Int in asexual blood stages of *P. falciparum*, cellular fractionation studies and western blot analysis were performed with rat anti-Pf-Int antibodies on *P. falciparum* cellular extracts. In general, the cytoplasmic fraction contains soluble proteins while the nuclear fraction contains nuclear, nuclear-associated, membrane-associated as well as intraorganellar, therefore also apicoplast proteins [32]. Fig 3A (top row lane 3) shows the characterization of Pf-Int in whole cell extract (wce), cytoplasmic (cyt) and nuclear (nuc) fractions for wild-type *P. falciparum* 3D7 ring stage parasites. The control shows a purified 35 kDa recombinant fragment of Pf-Int (C192) reaching from amino acid 192 to 490, the C-terminus [17]. A band of ∼60 kDa in size corresponding to the full length Pf-Int was identified in the nuclear fraction, therefore potentially associated with the apicoplast.

**Fig 3.**
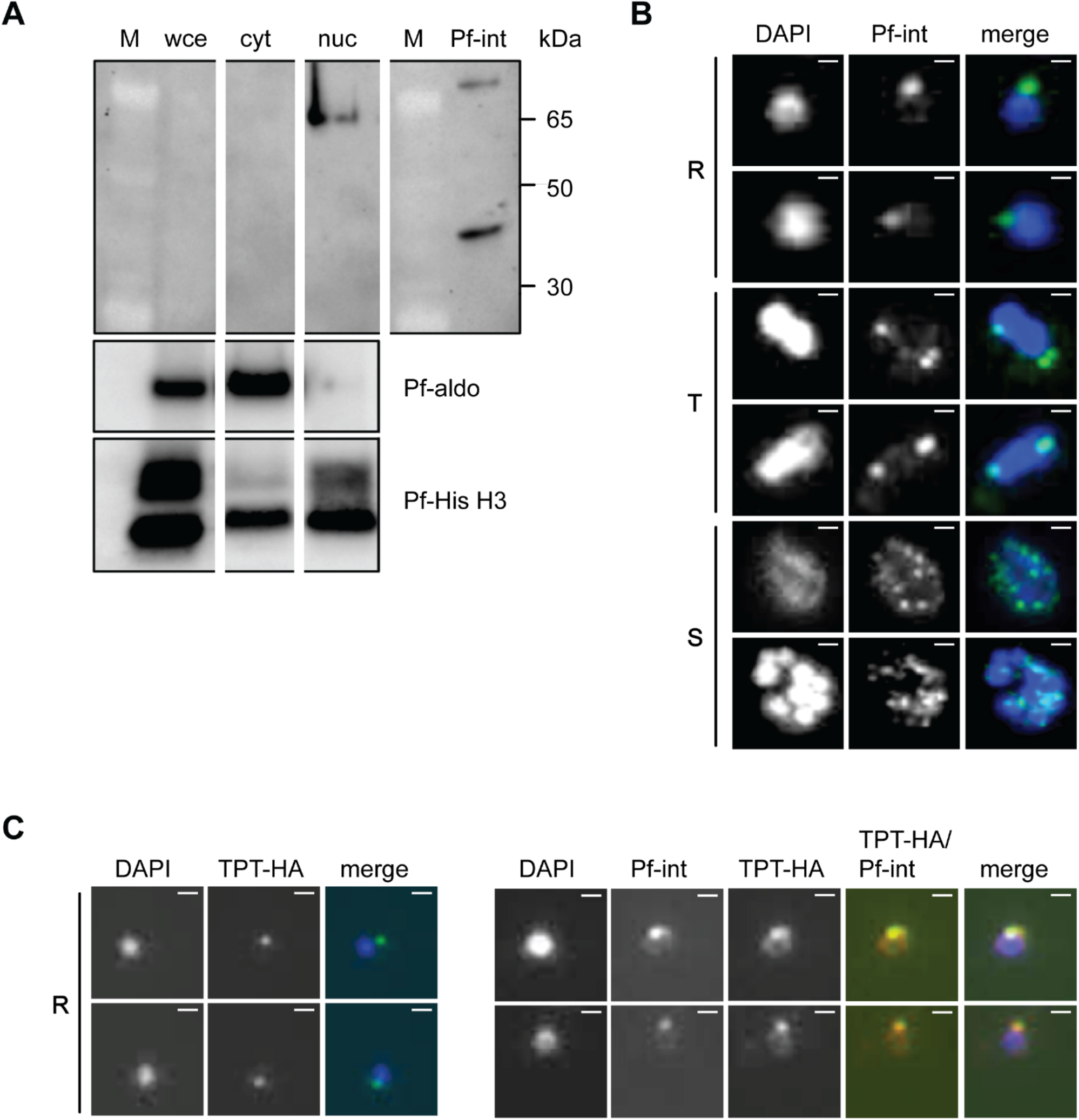
Subcellular localization of Pf-int. A) Pf-int localization studies by western blot. Detection of Pf-Int in synchronized P. falciparum 3D7 parasites: Ring stage whole cell extract (wce), cytoplasmic (cyt) and nuclear (nuc) fractions. rat-Anti-Pf-Int primary antibody dilution 1:500, secondary goat-anti-rat-HRP dilution 1:10,000. Developed with femto kit (ThermoFisher Scientific). Positive control: purified recombinant Pf-Int (C192, 35 kDa, 0.1 μg)(Top Row panel 4). Loading/ fractionation controls with cytoplasmic marker Pf-aldolase (Pf-aldo) and nuclear marker Pf-Histone H3 (Pf-His H3)(Lower panel). B) Initial characterization of anti-Pf-Int in immunolocalization studies. Anti-Pf-Int gave a punctate pattern, partly at the nuclear periphery (Panel 2 and 3). Shown for 3D7 ring (R), trophozoite (T) and schizont (S) stages. Anti-Pf-Int (1:250), secondary antibody AlexaFluor®488 (1:1000). The nuclei were stained with DAPI (Panel 1). Images are shown in duplicate for all parasite stages (Black vertical bar). C) Left panels: Anti-hemagglutinin (HA, green)-tagged PfoTPT localizes adjacent to the nucleus (blue) in PfoTPT-HA parasites. Right panel: Anti-Pf-Int (green) co-localizes with apicoplast-located PfoTPT (red) in PfoTPT-HA parasites. Each study is shown for ring (R) stages. Anti-Pf-Int primary antibody (1:250), mouse-anti-HA primary antibody (1:500); secondary antibodies AlexaFluor®488 (green) or AlexaFluor®568 (red) (1:1000). The nuclei were stained with DAPI. Images are shown in duplicate. Scale bar: 2μm.

Additionally, immunolocalization studies were performed using anti-Pf-Int antibodies on a mixed population of ring, trophozoite and schizont stage parasites of the wild-type 3D7 strain. Pf-Int was localized to a region adjacent to the nucleus, predominantly as a single punctum (Fig 3B, columns 2 and 3). This finding was in accordance with our results from the cellular fractionation studies. Based on these results, the co-localization of Pf-Int was investigated with the apicoplast marker outer-apicoplast-membrane triose phosphate transporter (PfoTPT) in TPT-HA transformed parasites [33]. The co-localization of Pf-Int with the apicoplast marker was confirmed (Fig 3C, columns 5, 7 and 8).

## Discussion

Overall, our protein localization studies suggest the presence of Int in an organelle adjacent to the nucleus, very likely the apicoplast. It may reach this organelle thanks to its bipartite leader peptide. Higher resolution studies such as electron microscopy would be needed to allow the exclusion of the possibility of a localization within the innermost algal-derived membrane of the organelle instead of the lumen (see Fig 2). In addition, the localization experiments of Int in Toxoplasma by Sheiner et al (2011) [26] have been reproduced and confirmed by Farhat and Hakimi (not shown). They observed that Int was co-localized with the apicoplast during different plastid segregation stages. Accordingly, it can be argued that Int may be intimately associated with DNA transactions, namely duplication of the plastid DNA, at least in *T. gondii*.

As a putative member of the Y-SSR family (conserved active site residues, see S1 Fig), Int could possibly play a role in apicoplast genome decatenation, or genome separation by hairpin resolution after rolling circle replication. Many of the replication machinery proteins in the apicoplast are known, but this is the first report of the presence of an enzyme with the function of a type I topoisomerase in the apicoplast. Nevertheless, it is noteworthy that Y-SSRs could fulfill the role of other enzymes such as DNA ligase or telomerase that may also be involved in the maintenance of apicoplast genome copy number. Both Pf-int and type I topoisomerase rely on the catalytic tyrosine present in the Y-SSRs for the cleavage of DNA and have significant similarities in sequence and structure [34, 35]. Topoisomerases proceed with either one or two strand cleavage (Type I/II), whilst Y-SSR usually cleave only one strand. Moreover, Y-SSRs require a specific sequence or structure, whereas topoisomerases are non-sequence specific [36]. The identification of specific DNA targets has so far been tricky, at least for Pf-Int. The potential DNA targets identified by SELEX and binding experiments by Ghorbal et al. (2012) [17] did not clearly yield a specific sequence motif. In this aspect, Pf-int could be similar to topoisomerases, which recognize DNA topology rather than specific sequences.

Our cellular fractionation experiments showed that Pf-Int is present in the nuclear (thus apicoplast-) associated fraction. Moreover, dividing cells, i.e. schizonts, also show Pf-int within each daughter parasite. Functioning as a topoisomerase would mean that Int could play a role in the actual machinery of the plastid DNA replication. Ligase and telomerase activities are usually needed just after replication in order to seal the endings of the newly synthesized DNA molecule. Knowing that the apicoplast genome is present in circular as well as in hairpin-closed linear form, generated by D-loop- or rolling circle replication [10], both types of functions would be needed in the apicoplast. HUH nucleases perform similar functions in rolling circle replication initiation and termination (Chandler et al 2013). HUH recognises hairpin structures (or the ssDNA at the 3’ or 5’ ends of the hairpin) [37] described to be present within the arms and at the center of the apicoplast genome’s IR. The possibility for Int to perform a HUH nuclease-like function is therefore also justified.

Additionally, we hypothesize that Pf-int could have a role in resolving genome concatemers that occur during rolling circle replication. If rolling circle replication had its origin in the centre region of the IR, this is where the resulting concatemer would have to be resolved in a site-specific manner. This scenario however, does not explain the existence of linear apicoplast DNA molecules. But such linear DNA could have their endings 5’-3’sealed by a telomerase-like enzyme similar to the hairpin telomere resolvase ResT in *Borrelia* [38]. No such enzyme has been described in the apicoplast as yet. HUH nucleases, have been reported to often have, apart from the HUH domain, domains with other activities, such as helicase or primase activity (Chandler et al. 2013). To this end, also a domain with telomerase function could be imagined. However, as similar as HUH nucleases and Y-SSRs may be, there are also striking differences: In contrast to Y-SSRs, which establish the phosphotyrosine at the 3’ end of the DNA, liberating a 5’ OH, HUH establishes the phosphotyrosine at the 5’ end, liberating the 3’ end for further processing (priming replication and/ or termination of replication). While Y-SSRs do not need high energy factors, HUH needs divalent metal ions for activity [37].

In this work, in line with the *T. gondii* integrase localization results of Sheiner et al. (2011) [26], we confirmed the probable sub-cellular localization of Pf-int to the apicoplast. This integrase, highly conserved among Apicomplexa, shown to be essential at least in *P. falciparum* [18], and localized to the apicoplast, where it may have an important function during DNA replication, could be a potential novel drug target against diseases caused by Apicomplexans. In the future, this study could be supplemented by pulling down Int together with apicoplast DNA, or atomic resolution Cryo-EM of Pf-Int bound to apicoplast, in order to confirm this enzyme’s function and validity as a drug target.

## Acknowledgments

Prof. Dr. Artur Scherf (Unité de Biologie des Interactions Hôte-Parasite, Institut Pasteur, Paris, F-75015) for kindly providing consumables and lab space to A. V. Berglar for IFA experiments.

Dr. Shuai Ding (Unité de Biologie des Interactions Hôte-Parasite, Institut Pasteur, Paris, F-75015; Prof. A. Scherf’s lab) for providing the *P. falciparum* TPT-HA parasite strain.

Dr. Dayana C. Farhat & Dr. Mohamed-Ali Hakimi (Institute for Advanced Biosciences (IAB), Team Host–Pathogen Interactions and Immunity to Infection, INSERM U1209, CNRS UMR5309, University Grenoble Alpes, Grenoble, F-38400) for testing the localization of Tg-Int in *T. gondii* and for contributing to the discussion in the manuscript.

## Author approvals

All authors have seen and approved the manuscript, and it hasn’t been accepted or published elsewhere.

