## Supplementary Material S1 and S2 for "Localization of the putative recombinase Pf-int to the apicoplast of *Plasmodium falciparum*"

### Supporting information

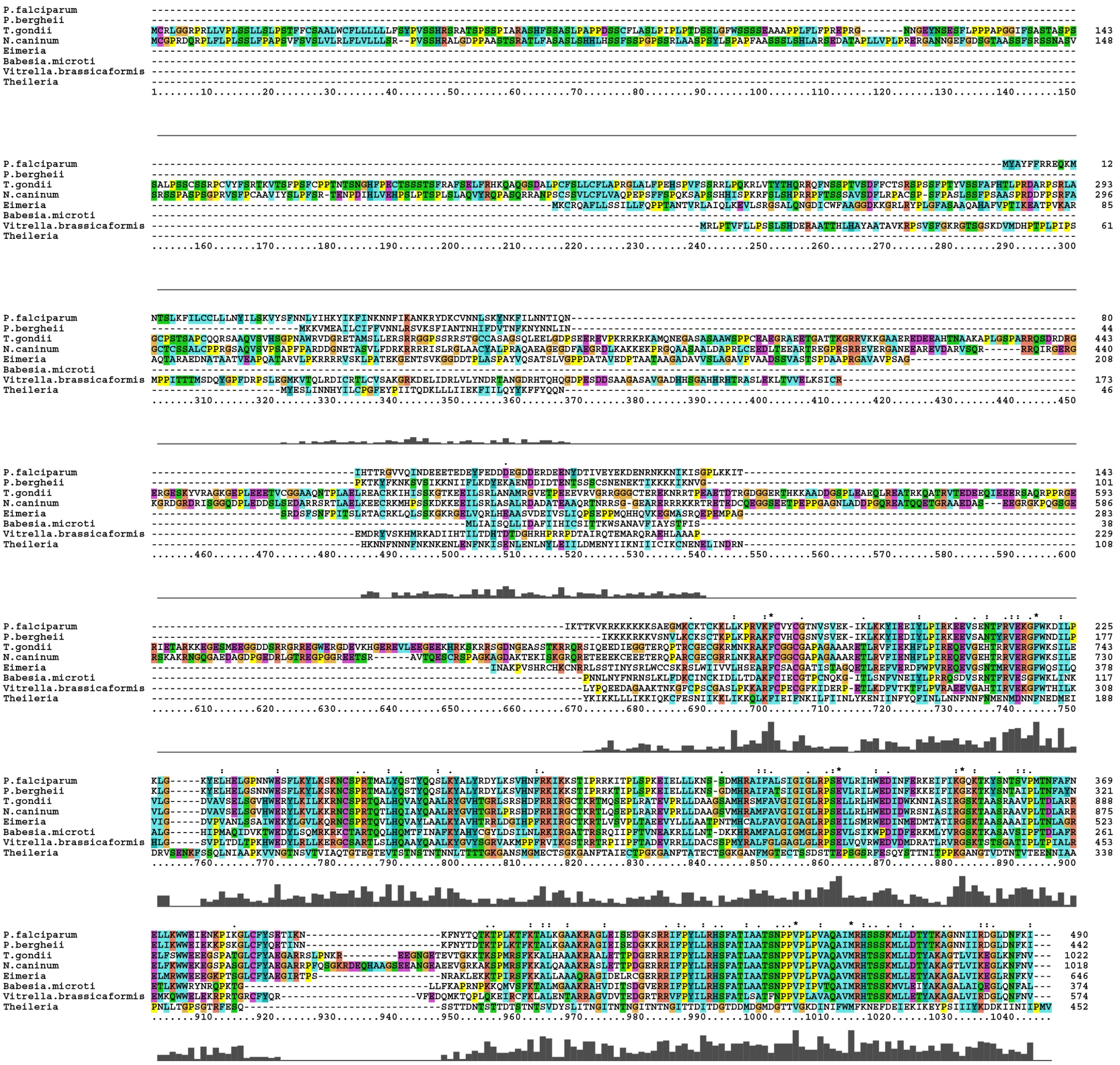


**S1 Fig. Apicomplexan Int putative protein sequence.** Alignment with ClustalX (Larkin *et al.*, 2007). Several members of the Int Tyr-recombinase protein were aligned to show homology using PAM % of average mutation table. In this alignment we compared: *P. falciparum* (XP_002809005.1), *P. berghei* (XP_034423895.1), *T. gondii* (XP_018637044.1), *N. caninum* (CEL66925.1), *E. maxima* (XP_013333234), *B. microti* (XP_012647521), *V. brassicaformis*

(CEM03382), *T. annulata* (XM_949126.1). Tyr-recombinases or related proteins were identified using BLAST (NCBI).

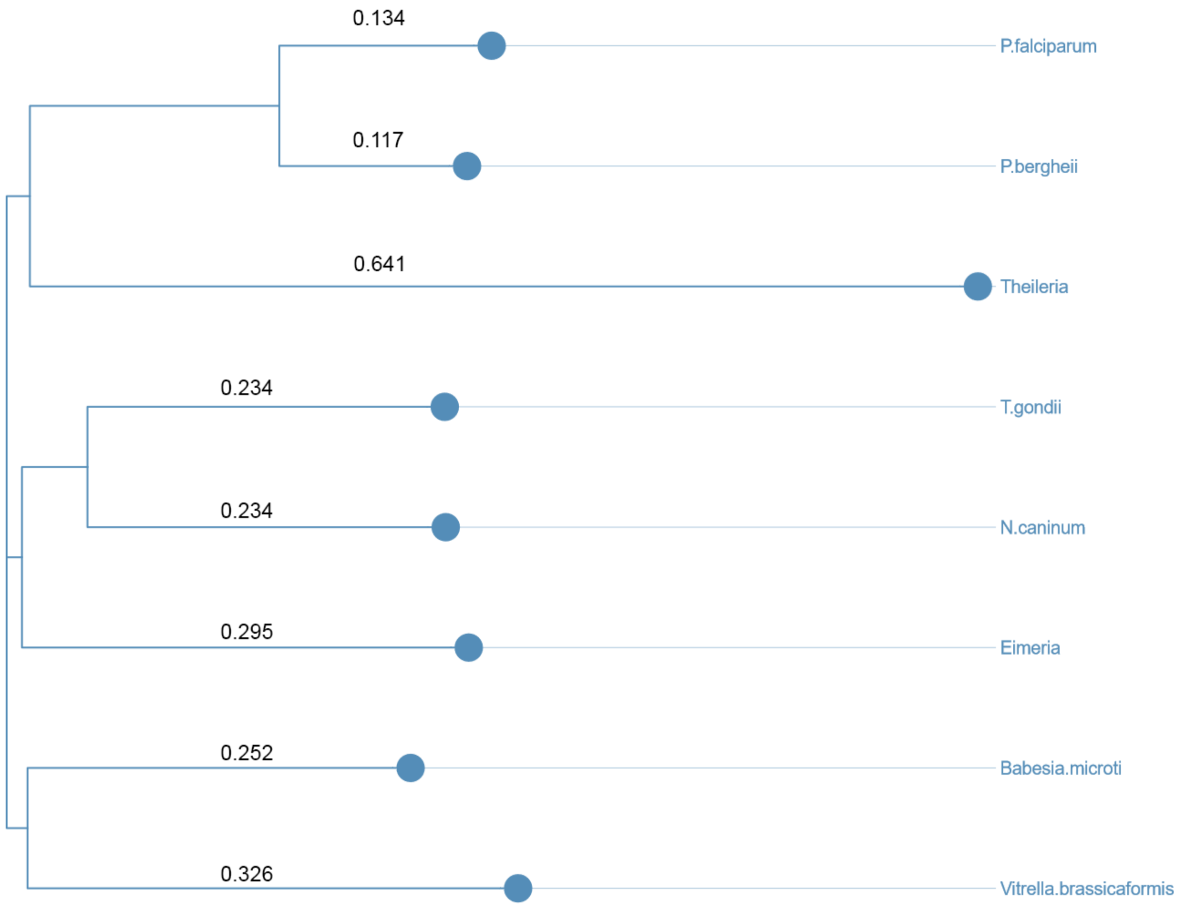


**S2 Fig. Phylogenetic tree of putative Tyr-recombinases from Apicomplexans.** The phylogenetic tree was computed using sequences from S1 Fig on the Galaxy Server (Institut Pasteur).
